# Overcoming host immune responses to an AAV-delivered HIV-1 bNAb in rhesus macaques mediated by co-delivery of PD-L1

**DOI:** 10.64898/2026.05.29.728806

**Authors:** Michael Kuipa, Abubakarr A. Koroma, Isai Leguizamo, Priya Dhole, Yash Barot, Michelle Y.-H. Lee, Gregory K. Tharp, Shan Liang, Magdalen Chouinard, Stephanie Ehnert, Stacey Weissman, Casey Whitehead, Rachelle L. Stammen, Jennifer S. Wood, Elizabeth H. Curran, Deepa Machiah, Evan D. Dessasau, Yoshiaki Nishimura, Jun Xie, Guangping Gao, Sumit Verma, Deanna A. Kulpa, Ian N. Moore, Steven E. Bosinger, Matthew R. Gardner

**Author notes:** Direct correspondence to: MRG.

## Abstract

Adeno-associated virus (AAV)-delivered anti-HIV-1 broadly neutralizing antibodies (bNAbs) have demonstrated promise for preventing and treating HIV-1 infection in preclinical models. However, host immune responses, specifically anti-drug antibodies (ADA), limit sustained bNAb expression. We have previously shown that PD-L1-mediated immune shielding improves the consistency of AAV-delivered bNAb 3BNC117 expression from muscle tissue in rhesus macaques. Here, we test the breadth of this approach with another bNAb, 10-1074. We show that AAV9.PD-L1 co-delivery with AAV9.10-1074 reduced the occurrence of ADA responses and improved the durability of bNAb expression for one year post administration. Notably 12 of 12 macaques that received AAV9.10-1074 vectors were protected against ten repeated SHIV_AD8-EO_ challenges. Histopathological profiling showed that AAV9.PD-L1 co-delivery prevented severe local inflammation and tertiary lymphoid structure formation at the administration site. Thus, immune shielding could serve as a broad strategy to prolong transgene expression from muscle-directed AAV-delivered biologics.

## Introduction

HIV-1 is a major global health challenge, with no vaccine or cure available to date. Antiretroviral therapy (ART) is highly effective at suppressing viremia and preventing new infections when taken prophylactically^1–4^, but requires lifelong daily adherence for most regimens^5^. Recently approved long-acting formulations lessen dosing frequency but share many of the limitations inherent to ART, including drug-related toxicities and viral rebound upon treatment interruption^6^. Immunotherapy with broadly neutralizing antibodies (bNAbs) could serve as an alternative to ART for HIV-1 therapy or prevention^7–11^. However, bNAbs are more costly to manufacture compared to ART and require repeated in-clinic administration multiple times a year to sustain antiviral efficacy, limiting the feasibility to meet the needs of both people living with HIV-1 or those at high risk of HIV-1 acquisition. Adeno-associated virus (AAV)-mediated delivery of bNAb genes to long-lived skeletal muscle cells addresses these limitations, offering the potential for years-long anti-HIV-1 immunity from a single administration^12^.

The prophylactic and therapeutic potential of single-dose AAV-delivered HIV-1 inhibitors has been demonstrated in humanized mice and nonhuman primates (NHPs)^13–17^. However, clinical translation remains limited by host immune responses, in particular the generation of anti-drug antibodies (ADA)^18–21^. To date, two clinical trials of AAV-delivered bNAbs have reported limited results, with low to undetectable bNAb concentrations and anti-bNAb ADA responses in participants^22,23^. Mitigating anti-bNAb immune responses is therefore a prerequisite for successful clinical translation. bNAbs vary in their immunogenicity based on their degree of somatic hypermutation (SHM), with higher SHM tending to correlate with higher anti-HIV-1 breadth and potency^24^. Notably, immune cell infiltration at AAV administration sites observed in both preclinical^25–27^ and clinical studies^22,28^ suggest that locally targeting the immune response at the site of transgene expression may be a viable strategy to enhance bNAb expression.

Current strategies to overcome immune responses against AAV-delivered transgenes have relied on systemic immunosuppression with calcineurin inhibitors^19^, mTOR inhibitors^29^, and corticosteroids^30^. These approaches are limited by cost, repeated in-clinic administration, drug toxicity, and increased infection susceptibility. Moreover, loss of benefit upon treatment withdrawal has been reported (citation needed). Alternative approaches to promote transgene tolerance include liver-targeted AAV delivery^31,32^ and exploiting the tolerogenic neonatal immune system through AAV-delivered bNAb administration at birth^33^.

We have previously shown that PD-L1-mediated immune shielding of transduced muscle enables durable expression of AAV-delivered bNAb 3BNC117 in rhesus macaques.

Additionally, PD-L1 co-delivery reduced ADA and rescued 3BNC117 protective efficacy against repeated simian (S)HIV challenges. PD-L1 signaling via PD-1, expressed on T cells, macrophages, and B cells following their activation is a well-characterized mechanism for limiting immune cell effector functions to promote immune tolerance and homeostasis^34^. In our study, PD-L1 co-delivery limited immune cell infiltration and consequently the development of tertiary lymphoid structures (TLSs) at the AAV administration site. TLSs are ectopic lymphoid organs that form at sites of chronic inflammation (including autoimmunity, cancer, and chronic infection) where they serve as privileged sites of antigen presentation and antibody production. Depending on context, TLSs can counteract pathology, as in cancer, or drive it, as in autoimmunity. Our findings implicate TLS formation at sites of transgene expression in skeletal muscle as a potential driver of the ADA response.

Beyond overcoming anti-transgene immune responses, optimized vector design is essential to maximize transgene expression to achieve prophylactic and therapeutic concentrations of AAV-delivered bNAbs. We have previously demonstrated the effects of capsid choice and transgene cassette expression elements in the absence of ADA on AAV-delivered antibody expression in head-to-head comparisons in rhesus macaques^35^. Specifically, using the AAV9 capsid and separation of bNAb heavy and light chain genes using a P2A ribosomal skipping peptide increased expression of a relatively low-SHM anti-SIV antibody. We applied these optimizations to our previous study with 3BNC117. Here, we investigated the breadth of our PD-L1 co-delivery strategy using the bNAb 10-1074, which has been tested in the clinic alongside 3BNC117 for HIV-1 therapy. Compared to 3BNC117, 10-1074 exhibits a lower degree of SHM, distinct HIV-Env epitope targeting (V3 glycan supersite vs CD4 binding site) and tends to exhibit lower immunogenicity in NHPs. Applying our vector optimization strategy alone showed that is was sufficient to sustain 10-1074 expression in 4 of 6 macaques, consistent with its relatively lower immunogenicity compared to 3BNC117. However, PD-L1 co-delivery virtually eliminated ADA leading to sustained expression in all 6 macaques. Additionally, PD-L1 co-delivery was associated with reduced inflammation and suppression of TLS formation at the site of AAV administration. Together with our previous work, these findings demonstrate the breadth of AAV9-vectored PD-L1 co-delivery as an immune shielding strategy applicable across multiple bNAbs.

## Results

### PD-L1 co-delivery improves the consistency of AAV-delivered 10-1074 expression

To assess whether PD-L1 co-delivery improves the durability of 10-1074 expression, we administered AAV9.10-1074 alone or co-delivered with AAV9.PD-L1 to two groups of macaques (n=6 per group) (**Fig. 1A**). AAV9 vectors were well tolerated and all macaques gained weight at the expected rate for the 52 week study (**Fig. S1**). One adverse event, anemia in macaque Mm029 at week 52, was reported and determined to be unrelated to AAV administration. We classified macaques as either sustained or transient bNAb expressors based on whether serum concentrations were maintained consistently ≥50 µg mL⁻¹. This threshold far exceeds concentrations required for protection in macaques^36^ and approaches concentrations considered necessary for therapy^10,11^. Transient expression, commonly observed in AAV-delivered bNAb studies, is characterized by an initial rise in serum bNAb concentrations followed by a sharp decline to low or undetectable amounts, coinciding with the emergence of ADA responses^18,20,21,26^. In the 10-1074-only group, 4 of 6 macaques achieved sustained expression (**Fig. 1B**). The remaining two (Mm019 and Mm022) had transient expression profiles associated with high ADA titers. Notably, Mm019 and Mm022 rebounded to concentrations ≥50 µg mL^-1^ by conclusion of the study, with ADA titers declining after week 30 and week 22 respectively, possibly reflecting tolerization. In contrast, PD-L1 co-delivery eliminated the transient phenotype entirely, with 6 of 6 macaques achieving sustained expression (**Fig. 1C**).

**Figure 1.**
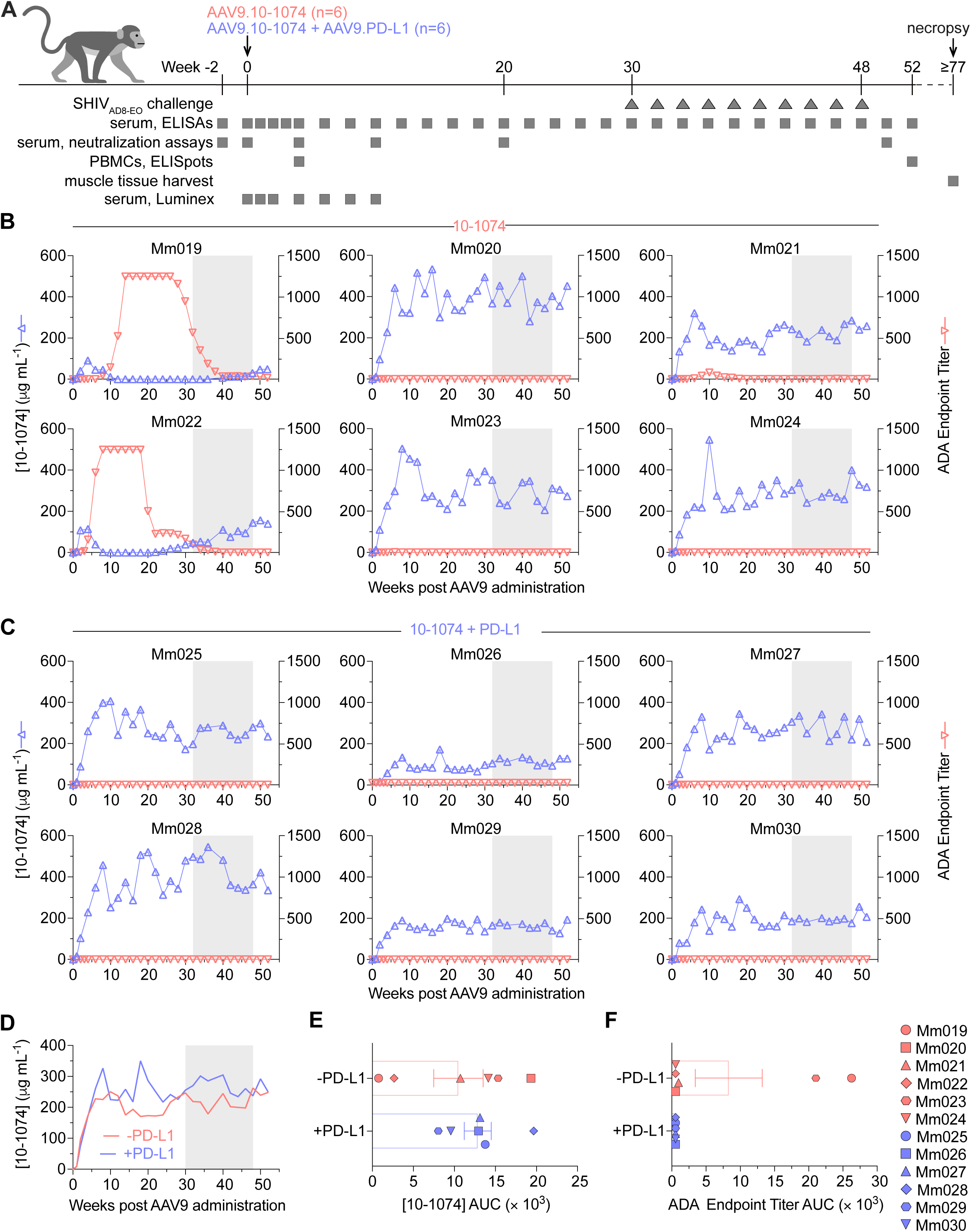
Longitudinal AAV9-delivered 10-1074 serum concentrations and ADA responses with and without PD-L1 co-delivery in rhesus macaques. **(A)** Schematic of the study. Two groups of rhesus macaques (n = 6 each) received AAV9.10-1074 alone or co-delivered with AAV9.PD-L1. Timeline indicates weeks post-AAV9 administration. Ten intrarectal SHIV_AD8-EO_ challenges were initiated at week 30. Samples were collected periodically for assays as indicated. Serum concentrations vs ADA endpoint titers in individual macaques that received **(B)** AAV9.10-1074 alone or **(C)** AAV9.10-1074 plus AAV9.PD-L1 as measured by gp120 or anti-10-1074 ELISA over 52 weeks. ADA endpoint titers are defined as the highest serum dilution with OD₄₅₀ ≥ 0.2. **(D)** Mean serum 10-1074 concentrations in macaques that received AAV9.10-1074. Gray shading in panels (B–D) indicates the SHIV_AD8-EO_ challenge phase. Comparison of mean AUC for **(E)** 10-1074 serum concentrations and **(F)** ADA endpoint titers for the 52-week study. Error bars represent SEM; symbols representing individual macaques are shared between (E) and (F).

Across both groups, macaques with sustained 10-1074 expression did not develop ADA responses beyond minor fluctuations above baseline. Average 10-1074 serum concentrations in individual macaques ranged from 15–351 µg mL⁻¹ in the 10-1074-only group vs 147–352 µg mL^-1^ in the 10-1074 plus PD-L1 group. Overall, average 10-1074 concentrations were similar between the two groups (**Fig 1D**) and no statistically significant differences in serum 10-1074 concentration or ADA endpoint titer area under the curve (AUC) were observed (**Fig. 1E,F).**

These results demonstrate that PD-L1 co-delivery promoted more consistent long term 10-1074 expression and underscore that ADA responses are an obstacle even for bNAbs with comparatively low immunogenicity.

### PD-L1 co-delivery does not reduce anti-AAV9 neutralizing antibodies or anti-10-1074 cellular immunity

While we observed the elimination of the host ADA response against 10-1074 in the co-delivered PD-L1 group, all 12 macaques developed neutralizing anti-AAV9 antibody responses after administration (**Fig. 2A,B**). There were no differences in anti-AAV9 antibody titer AUCs between groups (**Fig. 2C**), demonstrating that PD-L1 may only affect ADA against the expressed transgene. We then assessed anti-10-1074 cellular immunity using IFNγ ELISpot assays on week-4 and week-52 PBMCs (**Fig. 2D**). In both groups, we observed slight increases in cellular reactivity against all peptide pools tested by week 52. Notably, Mm027 in the 10-1074 plus PD-L1 group had the highest reactivity against all peptides. However, the peptide response did not appear to have an impact on the 10-1074 concentration once it reached its peak (**Fig. 1C**). Thus, while PD-L1 co-delivery markedly reduced anti-bNAb ADA responses, it did not consistently diminish anti-transgene peripheral cellular immune responses.

**Figure 2.**
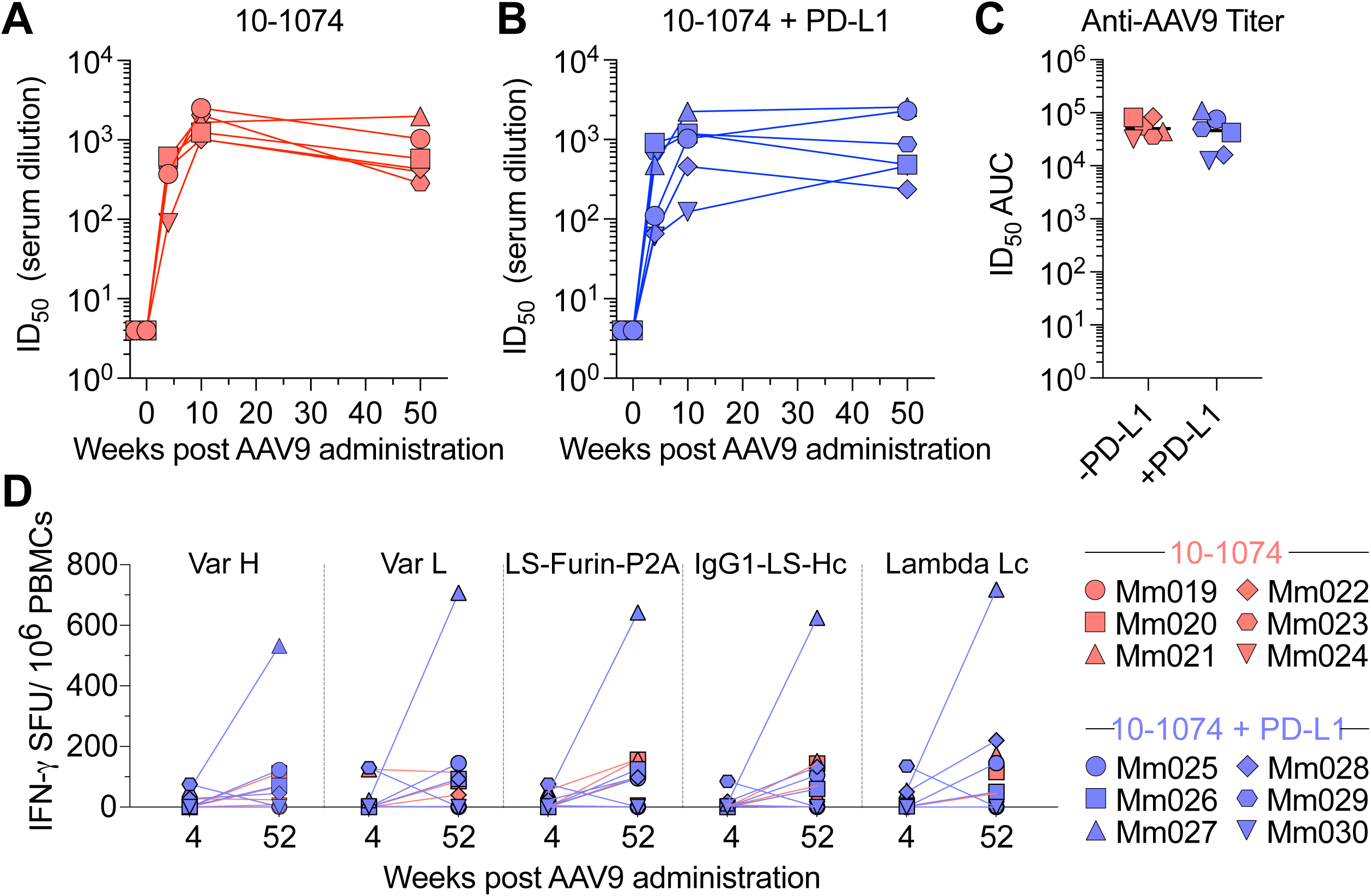
AAV9 neutralizing antibody responses and PBMC IFNγ ELISpot reactivity against AAV-expressed 10-1074. Serum AAV9 neutralization titers were measured at weeks-2, 0, 4, 10, and 50 using a HEK293T luciferase reporter neutralization assay. ID₅₀ values for each macaque are reported. Samples failing to reach 50% neutralization were normalized to a value <10. Week-2 serum served as baseline for each macaque. Serum ID₅₀ titers in individual macaques that received **(A)** AAV9.10-1074 only, or **(B)** AAV9.10-1074 plus AAV9.PD-L1. **(C)** ID_50_ AUC values for data in (A) and (B). Black lines indicate the median. **(D)** IFNγ ELISpot reactivity of PBMCs collected at weeks 4 and 52 against AAV9.10-1074 peptide pools (10-1074 VarH, 10-1074 VarL, LS-Furin-P2A peptide, IgG1 constant heavy chain region, and lambda light chain constant regions). Data expressed as spot forming units (SFU) per 10^6^ PBMCs. Note symbols representing individual macaques are shared in (A–D).

### AAV-expressed 10-1074 retains broad neutralization activity *ex vivo*

Unlike traditional vaccines, AAV-delivered bNAbs could offer predictable, broad and durable HIV-1 neutralization coverage from a single administration. To confirm that AAV-delivered 10-1074 retained functional neutralization activity in vivo, we assessed the serum neutralization activity in all macaques at 20 weeks post administration. At this timepoint, 10-1074 expression had stabilized. TZM-bl neutralization assays were performed using 11 HIV-1 pseudoviruses, simian-HIV (SHIV)_AD8-EO_ pseudovirus (Env of challenge virus), and SIVmac239 (negative control). As expected, macaques with measurable 10-1074 serum concentrations neutralized 10-1074-sensative isolates (**Fig. 3A,B**) and serum 10-1074 concentrations correlated strongly with SHIV_AD8-EO_ ID₅₀ neutralization titers across groups (p = 0.0001) (**Fig. 3C**).

**Figure 3.**
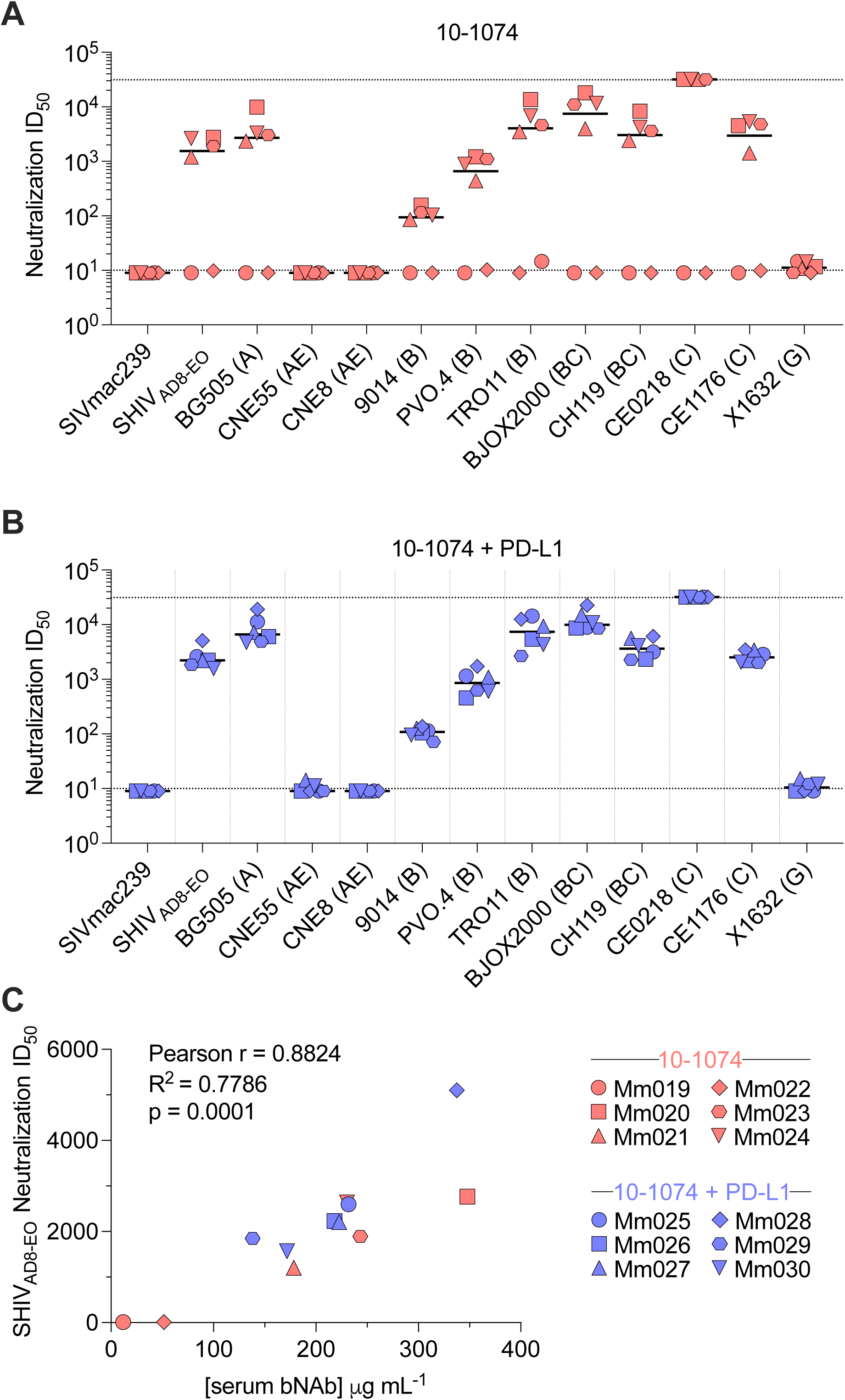
*S*erum HIV-1 neutralization breadth and potency following AAV9.10-1074 administration. Week-20 serum from all macaques was tested for neutralization against a panel of 11 HIV-1 pseudoviruses, the challenge virus SHIV_AD8-EO_ pseudovirus, and SIVmac239, using a TZM-bl neutralization assay. ID₅₀ values for each macaque per group are reported for the indicated pseudovirus and those samples that did not reach 50% neutralization were normalized to a value of <10. Week-20 serum ID₅₀ titers in individual macaques that received **(A)** AAV9.10-1074 only or **(B)** AAV9.10-1074 plus AAV9.PD-L1. In (A) and (B) HIV-1 clade is indicated in parentheses and black lines denote median ID₅₀ values. **(C)** Pearson correlation plot of serum 10-1074 concentration as determined in Fig. 1 and SHIV_AD8-EO_ ID₅₀ neutralization titer. Note symbols representing individual macaques are shared in (A–C). Pearson r value, R^2^ value, and p value (two-tailed t test), as determined by correlation analysis are shown. Statistical significance is defined as p value ≤ 0.05.

### AAV-delivered 10-1074 provides durable protection against repeated, intrarectal SHIV challenges

We have previously shown that PD-L1 co-delivery rescues the protective efficacy of AAV9-delivered 3BNC117 in intrarectal, low dose challenge experiments in rhesus macaques. We similarly evaluated the protective efficacy of AAV9.10-1074 in the current study, using the R5-tropic, tier-2 SHIV_AD8-EO_ strain. The 10 TCID₅₀ challenge dose, equivalent to 0.27 animal infectious doses (AID₅₀), was selected based on prior studies ^36^. At 30 weeks post-AAV administration, all macaques were challenged every two weeks for up to 10 total exposures or until infection was confirmed twice by qRT-PCR. All 12 macaques that received AAV9.10-1074 resisted 10 SHIV_AD8-EO_ challenges, which provided significant protection compared to our AAV9.PD-L1 historical control group (p < 0.0001) (**Fig. 4A,B**). During the challenge phase, average serum 10-1074 concentrations ranged from 9–398 µg mL^-1^ in the 10-1074-only group (**Fig. 1b**) and from 161–434 µg mL^-1^ in the 10-1074 plus PD-L1 group (**Fig. 1c**). Notably, Mm019 in the 10-1074-only group entered the challenge phase with undetectable serum 10-1074 concentrations. However, from week 36 onward (fourth challenge), Mm019 sustained concentrations above 0.16 µg mL⁻¹, the previously reported protective concentration for this SHIV_AD8-EO_ challenge stock^36^. In summary, AAV-delivered 10-1074 conferred complete protection against SHIV challenge irrespective of PD-L1 co-delivery, despite PD-L1 co-delivery improving the consistency of expression in macaques. This contrasts with our previous finding where PD-L1 co-delivery was necessary to rescue protection mediated by AAV-delivered 3BNC117. Taken together, these findings establish transgene immunogenicity as a factor of AAV-vectored bNAb in vivo function, with PD-L1 co-delivery serving as a practical safeguard against loss of prophylactic efficacy.

**Figure 4.**
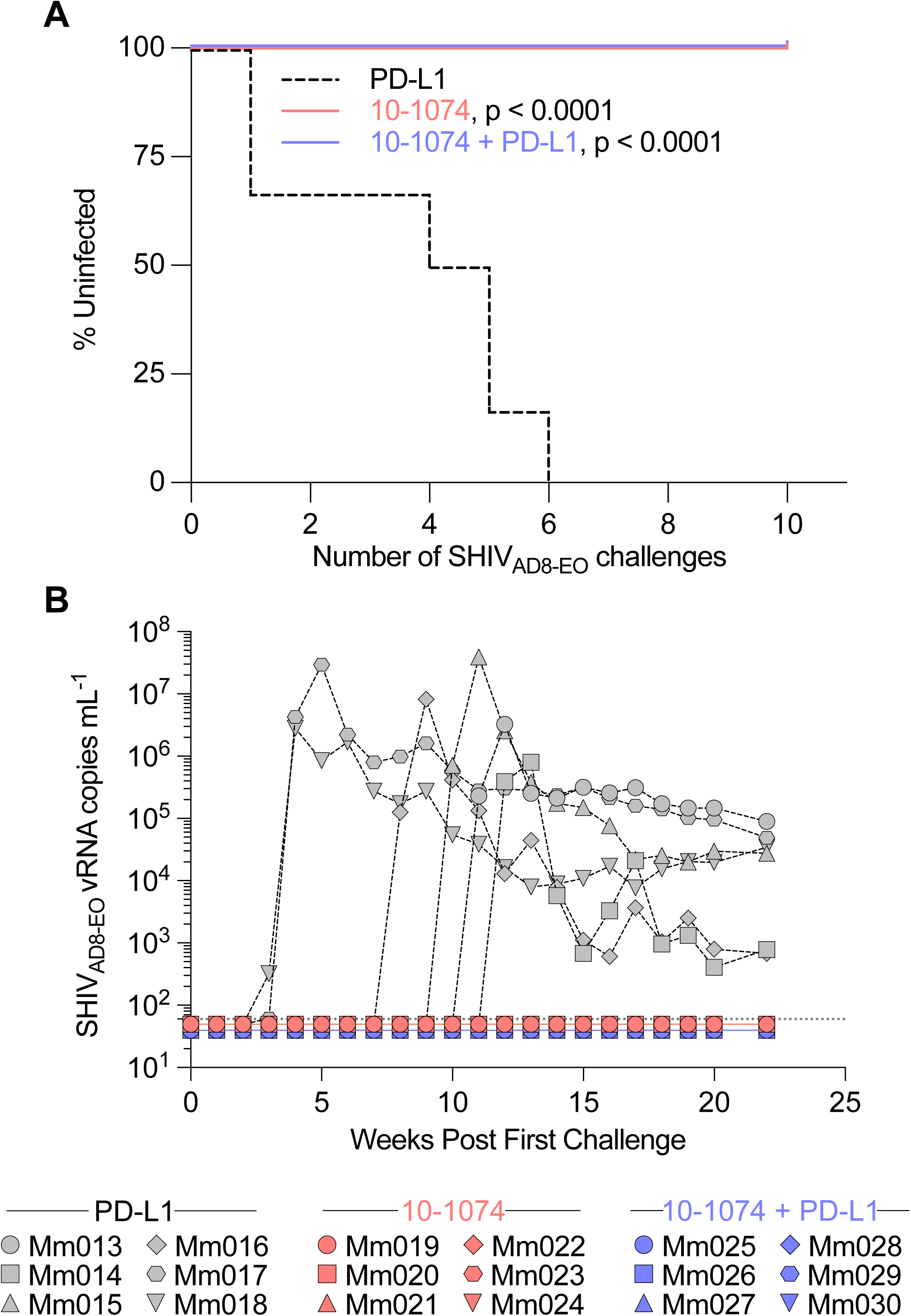
AAV9.10-1074-mediated protection against intrarectal SHIV_AD8-EO_ challenges. **(A)** Kaplan-Meier analysis of protection from ten biweekly intrarectal SHIV_AD8-EO_ challenges infection in macaques that received AAV9.10-1074 with or without PD-L1 co-delivery. A historical cohort of macaques (n = 6) that received AAV9.PD-L1 alone served as the control group. **(B)** Plasma SHIV_AD8-EO_ viral RNA mL^-1^ levels during the SHIV_AD8-EO_ challenge phase over time in macaques from (A). Viral loads were measured by qRT-PCR with a limit of detection of 60 copies mL^-1^, indicated by the dotted line. Statistical significance was determined by Mantel-Cox test.

### PD-L1 co-delivery lessens inflammation severity at the AAV administration site

We have previously shown that the degree of inflammation at the AAV administration site correlates with ADA severity. Histological examination of hematoxylin and eosin (H&E)-stained quadriceps muscle sections collected near the site of AAV administration from all 12 macaques in the current study revealed a similar association (**Fig. 5A**). Inflammation severity was scored blindly on a five tier scoring system: “-“, “+/-”, “+”, “++” or “+++”. For reference, previously reported macaques receiving AAV9.PD-L1 alone had inflammation severity scores ranging from “-“ to “+”, with a median of “+/-”, likely reflecting AAV capsid-driven inflammation. 10-1074-only group macaque Mm019, which had the highest ADA endpoint titer AUC, also had the highest inflammation severity score (“++”) (**Fig 5A, B**). We observed a similar distribution of inflammation severity scores in 10-1074 plus PD-L1 group macaques to the PD-L1-only historical controls, with a range of “-” to “+” and a median of “+” (**Fig. 5A, C**). Thus, in agreement with our previous study, we find that inflammation at the AAV administration site serves as a proxy for the magnitude of the ADA response.

**Figure 5.**
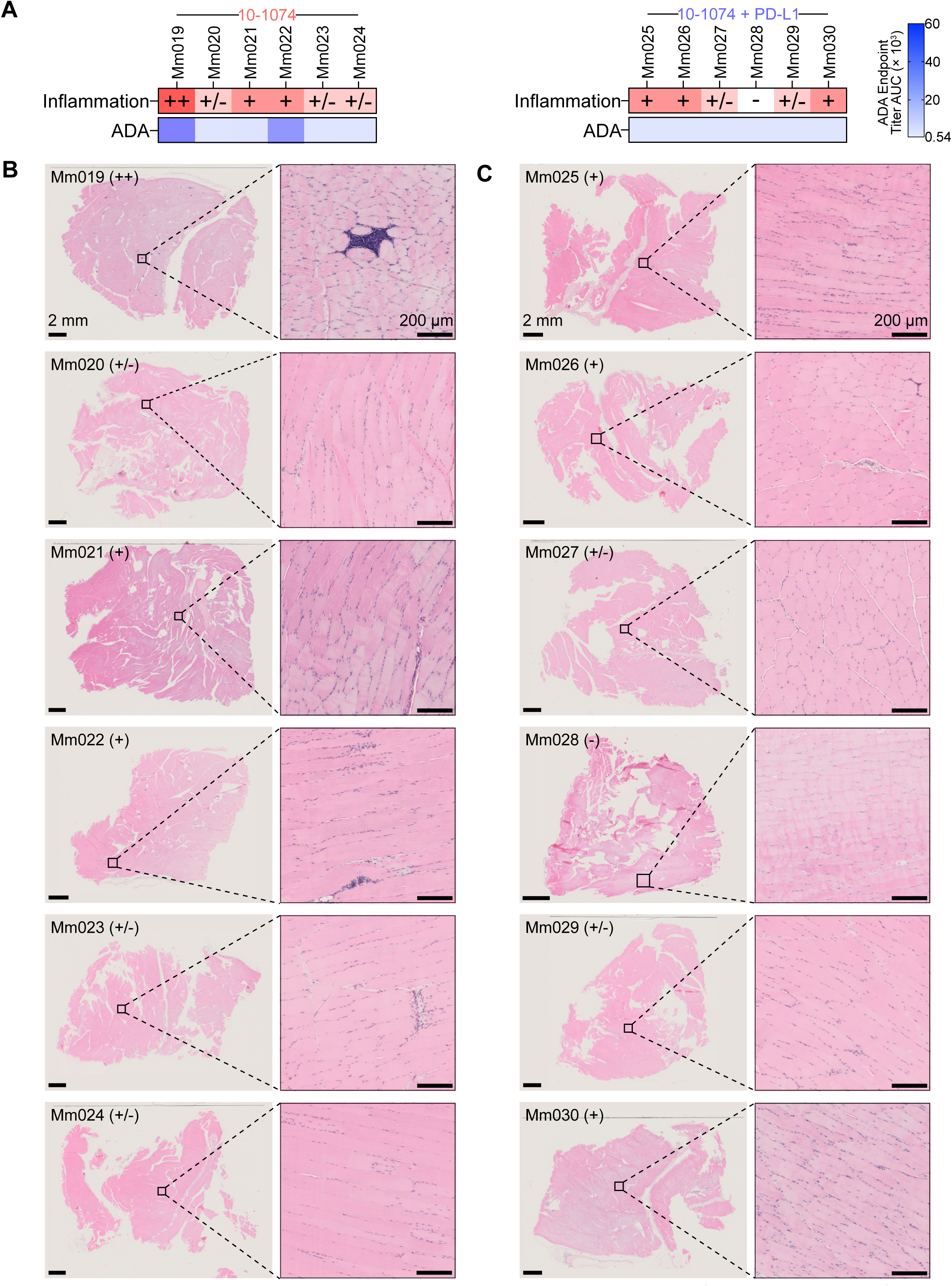
Histological assessment of muscle tissue collected near the site of AAV9.10-1074 administration. Upper left quadriceps muscle tissue was harvested at necropsy from all 12 macaques and H&E-stained sections were scored for inflammation. **(A)** Muscle inflammation severity scores vs ADA endpoint titer AUC determined in (Fig. 1) for all 12 macaques. Darker colors indicate higher inflammation (red) or ADA endpoint titer (blue). Inflammation scoring legend:-, no evidence of interstitial hypercellularity or overt inflammation; +/-, slight interstitial hypercellularity or minimal focal inflammatory cell infiltration; +, moderate interstitial hypercellularity or mild, diffuse inflammation; ++, evident inflammation with small-to-moderate foci showing multifocal distribution; +++, marked inflammation with moderate-to-large foci displaying multifocal and coalescing distribution. Severity scores were adjusted to accommodate changes to normal muscle fiber morphology such as muscle degeneration and/or necrosis. H&E stained tissue in individual macaques that received **(B)** AAV9.10-1074 alone or **(C)** AAV9.10-1074 plus AAV9.PD-L1. Scale bars are 2 mm on the left and 200 µM on the right.

### TLSs at the AAV administration site are associated with ADA responses

A key finding of our previous study was that TLSs form in the muscle at the site of AAV administration and are associated with the magnitude of the ADA response. Consistent with this, immunohistochemical analysis of quadriceps muscle sections from Mm019, which had the highest ADA endpoint titers in the current study, revealed evidence of TLS formation. We stained for the markers CD3, CD4, Foxp3, CD56, CD19, CD68, and PD-1 (**Fig. 6A**). In line with our previous study, we identified T cells, NK cells, B cells, and macrophages. We observed several germinal center-like regions with segregated CD4 and CD19 staining (**Fig. 6B**). We also detected PD-1 and FoxP3 expression within the germinal center-like structures, potentially indicating the presence of T follicular regulatory (Tfr) cells, T follicular helper (Tfh) cells, and/or T regulatory (Treg) cells. To investigate the kinetics of TLS formation in Mm019, we examined the serum concentrations of TLS-associated cytokines and chemokines CXCL13, IL-1β, IL-6, IL-7, IL-13, IL-13, IL-17, IL-21 and TNF-α by Luminex over the first 10 weeks following AAV administration (**Fig. 6C**). We did not detect significant changes in the concentrations of any of the analytes tested. This suggests that TLS-associated inflammatory signaling may be local and require profiling at the administration site to determine the kinetics of TLS formation.

**Figure 6.**
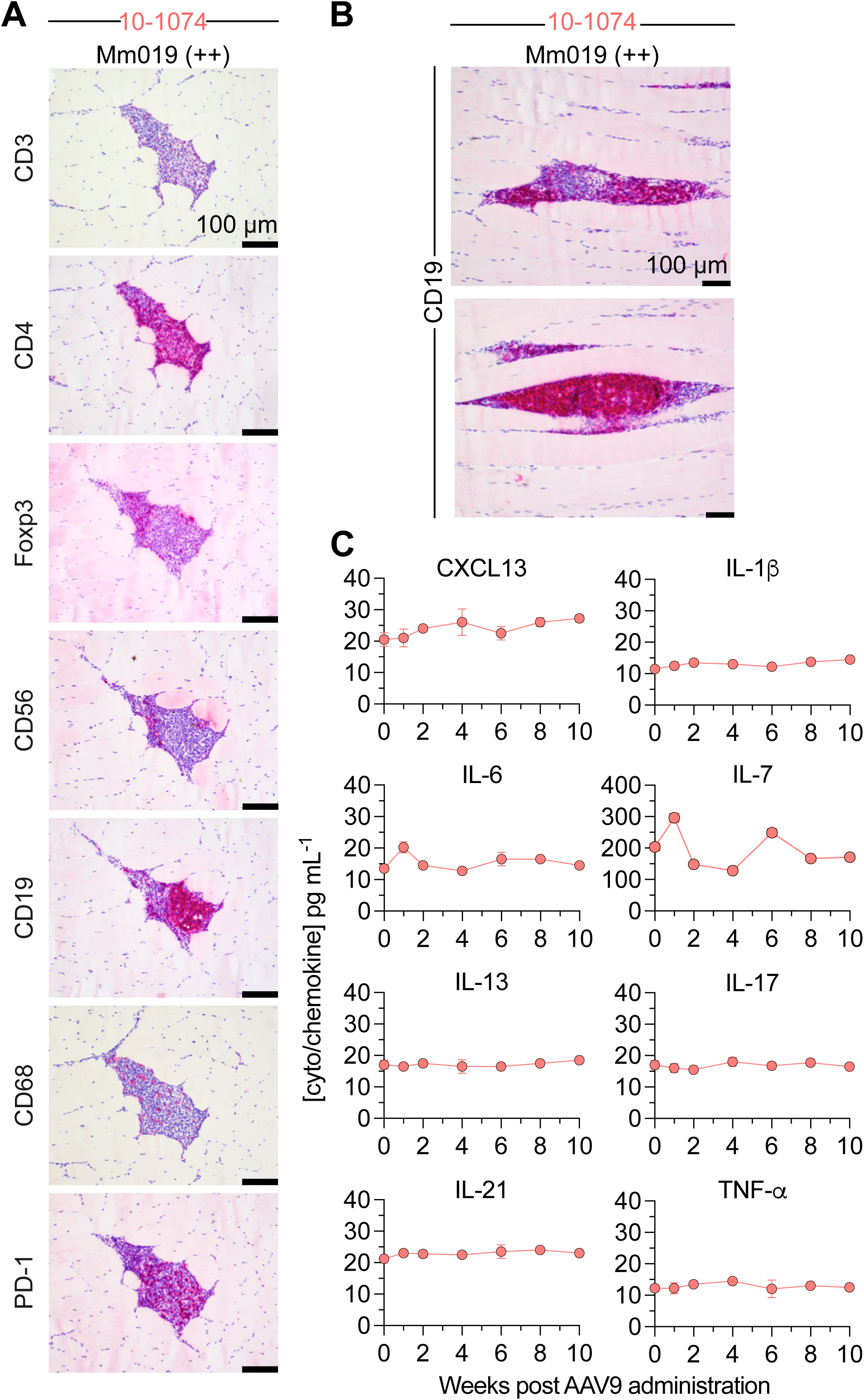
TLS formation at the site of AAV9.10-1074 administration and serum cytokine profiling in a macaque with high ADA response. **(A)** IHC staining for CD3, CD4, Foxp3, CD56, CD19, CD68, and PD-1 in Mm001. (**B**) IHC staining for CD19 in Mm001 showing TLSs. (**C**) Serum cytokine and chemokine levels in Mm001 over the first 10 weeks post-AAV9.10-1074 administration, quantified by Luminex. Scale bars in (A) and (B) are 100 µM.

Nevertheless, the presence of TLSs exclusively in the macaque with the highest ADA magnitude suggests that TLSs may contribute to the magnitude of the ADA response.

## Discussion

Here we evaluated the breadth of our AAV.PD-L1 approach to improve bNAb expression in rhesus macaques through reducing host immune responses. PD-L1 co-delivery prevented detectable anti-10-1074 ADA responses, allowing for sustained 10-1074 expression (≥50 μg mL⁻¹) in 6 of 6 macaques. In contrast, 4 of 6 macaques in the 10-1074-only group achieved sustained expression with two macaques developing anti-10-1074 ADA responses. Our previous work delivering AAV1.10-1074 vectors in rhesus macaques prior to our vector optimization strategies yielded concentrations <5 μg mL⁻¹ in 6 of 6 macaques. Thus, our optimizations of the capsid, promoter, and 2A selections alone resulted in dramatic improvements in 10-1074 expression that was further augmented through co-delivered PD-L1. Additionally, these findings extend our previous results with 3BNC117, suggesting that PD-L1 co-delivery may be a broadly applicable strategy for maintaining the expression of AAV-delivered bNAbs.

The 10-1074 expression levels achieved here represent, to our knowledge, the highest AAV-delivered bNAb concentrations reported consistently across multiple rhesus macaques. In the best-performing prior study, AAV9-delivered 10-1074 reached 5–55 μg mL⁻¹ in 5 macaques over the majority of a 56-week study and required an intensive 14-week rapamycin regimen. These concentrations achieved here have been shown to be above those concentrations needing to suppress an HIV-1 infection without ART. These results suggest some flexibility to incorporate optimizations aimed at improving safety (which may reduce expression) such as the use of muscle-specific promoters and AAV-dose reduction.

Consistent with our previous findings, high anti-bNAb ADA responses were again accompanied by TLS formation at the AAV administration site. Across both studies, 5 of 7 macaques that developed high anti-bNAb ADA endpoint titers also presented with TLSs. Importantly, we have not observed TLSs in any of the 18 macaques receiving PD-L1 across the two studies we have performed. Given that ADA responses typically peak at 6–8 weeks post-AAV administration, we reasoned that TLSs were forming early but did not detect changes in TLS-associated cytokines and chemokines in the one animal presenting with TLSs in this study. Our inability to find a marker of TLS development in the blood indicates that in situ analysis of the administration site is required to capture the early immune dynamics driving TLS formation.

We observed complete protection from repeated intrarectal SHIV_AD8-EO_ challenges in all macaques. This contrasts with our previous study, where PD-L1 was necessary to rescue the protective efficacy of 3BNC117. However, complete protection against a single SHIV strain does not capture the breadth of coverage that would be required in a real-world setting, where circulating HIV-1 strains vary in neutralization sensitivity. The Antibody Mediated Prevention (AMP) trials^9^ established that bNAb VRC01-mediated protection was concentration-and neutralization sensitivity-dependent, with prevention efficacy of ∼75% against viruses with IC_80_ values <1 μg mL⁻¹, but approaching zero for viruses with IC_80_ values >1 μg mL⁻¹. PT_80_, defined as the ratio of serum bNAb concentration to the in vitro IC_80_ of the target virus, has been proposed as a predictive biomarker of bNAb prevention efficacy, with PT_80_ >200 associated with ∼90% prevention efficacy in the AMP trials. In our study the lowest individual mean 10-1074 concentration was 15 µg/mL without PD-L1 co-delivery and 147 µg/mL with PD-L1 co-delivery, representing an approximately 10-fold increase. This proportionally expands the IC80 threshold below which PT_80_ >200 is maintained, from 0.075 µg/mL to 0.74 µg/mL, predicting more consistent protection against a broader range of circulating viral strains.

PD-L1 co-delivery reduced ADA responses against the bNAb transgene but did not attenuate neutralizing antibody responses against AAV9 capsid, suggesting that redosing with the same serotype would remain limited with our approach. This is not surprising; upon AAV9 administration, APCs immediately encounter a high concentration of antigen and can initiate an immune response before PD-L1 expression has occurred. In contrast, the transgene accumulates gradually, such that APCs encounter antigen in the presence of PD-L1, enabling local suppression of T cell responses and limiting downstream ADA development. Furthermore, pre-existing cross-reactive memory B cells and the dense, repetitive array of B cell epitopes across the ∼60 AAV9 VP monomers may drive T cell-independent humoral responses against the capsid, which are features less susceptible to PD-L1-mediated local immune shielding and not applicable to the secreted bNAb.

Several rodent studies have demonstrated the capacity of AAV-delivered PD-L1 to improve the consistency of transgene expression^26,37–39^. A foundational study showed that AAV1-delivered PD-L1 or PD-L2 combined with CTLA-4-Ig improved tolerance to AAV1-delivered ovalbumin in C57BL/6 mice, although PD-L1 or PD-L2 delivery alone did not^26^. Notably, they concluded that PD-L1 or PD-L2 cannot interfere with systemic immune priming and instead regulate T cell effector functions locally. This is consistent with our observation of similar IFNγ ELISpot anti-transgene PBMC reactivity, regardless of PD-L1 co-delivery, as was observed in our previous 3BNC117 study. In their expression cassettes, PD-L1 was driven by the slower CBA-based core promoter while antigen was under the faster CMV promoter, potentially delaying immune shielding relative to antigen expression. We reasoned that earlier PD-L1 activity would be required and reversed this configuration in our approach. Subsequent studies have shown that PD-L1 co-delivery reduced loss of AAV9-delivered luciferase expression following IV delivery and enhanced AAV6-delivered muSEAP expression in AAV6-pre-immunized but not naïve mice following IM injection^39^. In a rat lung transplant model, AAV9-delivered PD-L1 combined with abatacept also attenuated acute cellular rejection severity^38^. Our findings validate PD-L1-mediated immune shielding across a second bNAb in an NHP model, further supporting its clinical applicability for AAV-delivered biologics. Broadly, this work reinforces the view that the local immune environment at the AAV administration site is the principal determinant of ADA development in the context of IM delivery, with TLS formation as a key underlying mechanism.

Beyond its translational implications, the ability of PD-L1 co-delivery to suppress ADA responses in NHPs offers practical utility for evaluating vector design and transgene performance in preclinical studies. In the accompanying study by Leguizamo et al., for example, we leverage PD-L1 co-delivery to attenuate ADA responses and more reliably assess how promoter selection within AAV transgene cassettes influences 10-1074 expression in rhesus macaques. The present study therefore establishes PD-L1 co-delivery not only as a strategy for enhancing bNAb expression, but as a practical preclinical platform for gene therapy development involving highly immunogenic transgenes.

## Materials and Methods

### Rhesus macaques

Twelve Indian-origin rhesus macaques (*Macaca mulatta*) housed at the Emory National Primate Research Center (ENPRC) in Atlanta, Georgia were enrolled in this study. The cohort comprised 9 males and 3 females aged 3.8–9.7 years and weighing 6.3–11.1 kg at the time of vector administration. Macaques were stratified into two groups based on age, weight, sex, and baseline serum AAV9 neutralizing antibody (NAb) titers (all ID_50_ titers were <1:10). All macaques were SIV-negative and *Mamu-B*08*⁻/*Mamu-B*17*⁻. Macaque identifiers, *Mamu-A*01* and *Mamu-A*02* genotype, sex, weight, and age at study initiation are detailed in **Table S1.** Macaques were housed in pairs with compatible animals for the duration of the study. All animal procedures were conducted in accordance with the National Institutes of Health (NIH) Guide for the Care and Use of Laboratory Animals (8th Edition), the United States Department of Agriculture (USDA) Animal Welfare Act, and institutional guidelines, under protocols approved by the Emory University Institutional Animal Care and Use Committee (Permit No. 202100453). The animal care facilities at ENPRC are accredited by the USDA and AAALAC, International.

### AAV Production and Purification

AAV9.10-1074 and AAV9.PD-L1 transgenes were synthesized and codon optimized by GenScript. Transgenes were cloned into an AAV transfer plasmid including the AAV serotype 2 inverted terminal repeats (AAV2-ITRs) using NotI restriction sites. AAV transgene expression cassettes have been previously described. Briefly, the 10-1074-encoding cassette comprised a CMV enhancer, chicken beta-actin core promoter, and SV40 intron driving expression of 10-1074 heavy and light chains connected via a furin cleavage site (RKRR), SGSG linker, and P2A ribosomal skipping peptide. Downstream regulatory elements included a woodchuck hepatitis virus posttranscriptional regulatory element and SV40 polyadenylation signal. The rhesus macaque IgG1 constant heavy chain region incorporates M428L/N434S half-life-extending substitutions. The PD-L1 cassette comprised a CMV enhancer, CMV promoter, SV40 intron, PD-L1 coding sequence, WPRE, and SV40 polyadenylation signal. Recombinant AAV was produced by the University of Massachusetts Medical School Vector Core and has been previously described^40^. Briefly, HEK293T cells were transfected with either 10-1074- or PD-L1-encoding transfer plasmids, a plasmid encoding AAV2 rep and the AAV9 capsid, and a helper plasmid encoding adenovirus genes. After collection of transfected cell lysates, AAV9 vectors were purified using three consecutive CsCl centrifugation steps. Vector genome (vg) copy number was determined by qPCR. AAV particle quality and purity were assessed by electron microscopy and silver-stained SDS-PAGE.

### AAV vector administration

AAV9.PD-L1 and/or AAV9.10-1074 were administered intramuscularly across eight sites (two injections in each of the lower and upper quadriceps, biceps, and deltoid muscles) at a dose of 2.5 × 10¹² vg kg⁻¹ per vector. Vectors were diluted in sterile PBS at a volume <1 mL.

### Protein Production and Purification

HEK293T cells (ATCC) were cultured in DMEM supplemented with 10% fetal bovine serum (FBS), 0.5% penicillin-streptomycin, and 10 mM HEPES, and maintained at 37°C with 5% CO₂. Recombinant 10-1074 was produced by co-transfecting HEK293T cells in T225 flasks with an AAV transfer plasmid encoding 10-1074 and a furin expression plasmid using PEIpro transfection reagent (Polyplus) according to the manufacturer’s instructions. The expression plasmid encoding furin has been previously described^21^. After 24 h, DMEM was replaced with FreeStyle 293 Expression Medium (Invitrogen). Media collected 48 h later was then clarified by centrifugation at 4,500 × g for 10 min and filtered through 0.45 µm filter flasks (Thermo Scientific). Antibody was purified by protein A affinity chromatography using HiTrap MabSelect SuRe columns (Cytiva) and eluted with IgG Elution Buffer (Thermo Scientific) into 1 M Tris-HCl (pH 8). Eluate was then buffer exchanged into PBS using Amicon Ultra centrifugal filter tubes (Sigma). Antibody heavy and light chain composition was assessed by Coomassie-stained SDS-PAGE

### Challenge virus production and challenge procedure

Preparation of the rhesus macaque PBMC-derived R5-tropic, tier-2 SHIV_AD8-EO_ stock has been previously described^41^. Challenge virus was diluted in 1.0 mL of serum-free RPMI medium to 10 TCID_50_, after which it was loaded into a 3 mL syringe and delivered intrarectally to each macaque. The 10 TCID₅₀ challenge dose is equivalent to 0.27 animal infectious doses (AID₅₀) and was selected based on prior studies^36^. Beginning at 30-weeks post AAV administration, macaques were challenged every two weeks for up to 10 total exposures or until infection was confirmed twice by qRT-PCR. The SHIV_AD8-EO_ molecular clone was a gift from Dr. Malcolm Martin.

### SHIVgag Plasma Viral Load Quantification

Quantification of SHIV plasma viral load has been previously described^42^. Briefly, RNA was extracted from macaque plasma using the QIAsymphony DSP Virus/Pathogen Mini Kit (Qiagen), according to the manufacturer’s recommendations, with a 200 µL input volume and 60 µL elution volume. SHIV RNA was quantified by one-step qRT-PCR using the TaqMan Fast Virus 1-Step Master Mix (Applied Biosystems) and SHIV/SIVgag-specific primers and probe (forward: 5′- GCAGAGGAGGAAATTACCCAGTAC-3′, Fisher Scientific; reverse: 5′-CAATTTTACCCAGGCATTTAATGTT-3′, Fisher Scientific; probe: 5′-6FAM-TGTCCACCTGCCATTAAGCCCGA-TAMRA-3′, Applied Biosystems). A standard curve was generated from five-fold serial dilutions of SHIV/SIVgag plasmid RNA ranging from 2.11 × 10^8^ to 2.69 × 10^3^ copies mL^-1^, which were run in duplicate on each plate. Reactions were performed with the following cycling profile: 50°C for 15 min, 95°C for 2 min, 40 cycles of 95°C for 15 s, and 60°C for 1 min on the Applied Biosystems 7500 or QuantStudio 3 Real-Time PCR System. SHIV RNA concentrations (copies mL^-1^) of plasma were calculated from the standard curve (R² ≥ 0.99) and adjusted for sample dilution if applicable. The assay limit of detection is 60 copies mL^-1^.

### TZM-bl Neutralization assay

HIV-1, SHIV, and SIV pseudoviruses were produced as previously described^15,35,43–45^. Briefly, heat inactivated sera were diluted 1:5 in DMEM and then serially diluted five-fold, yielding final assay dilutions from 1:10 to 1:31,250. Each dilution was tested in duplicate in a 96-well plate. Diluted sera were then mixed 1:1 (v/v) with pseudoviruses in DMEM and incubated at 37°C for 30 min. Next, 10,000 TZM-bl cells (NIH AIDS Reagent Program; HRP-8129; contributed by Drs. John C. Kappes and Xiaoyun Wu, and Tanzyme, Inc.) were added to each well. After incubation at 37°C for 48 h, luciferase activity was determined using Britelite Plus (Revvity) and read on a BioTek Synergy Neo2 plate reader (Agilent). ID₅₀ values were determined by fitting the data to a four-parameter logistic regression model. Expression plasmids used for pseudovirus production for pNL4-3Δenv, BG505, and SIVmac239 have been previously described^43,44^. The following reagents were sourced from the NIH AIDS Reagent Program: TRO11, CNE8, BJOX2000, X1632, CE1176, CH119, CE0217, and CNE55 (cat# 12670; contributed by Dr. David Montefiori); PVO.4 (ARP-11022; contributed by Drs. David Montefiori, Feng Gao, and Ming Li); and 9014 (ARP-11571; contributed by Drs. Beatrice H. Hahn, Brandon F. Keele, and George M. Shaw).

### gp120 and ADA ELISAs

To quantify serum 10-1074 concentrations, Costar 96-well half-area assay plates were coated with 3 µg mL^-1^ gp120-ADA (Immune Technology) in PBS overnight at 4°C. For ADA ELISAs, assay plates were coated with 3 µg mL^-1^ of recombinant 10-1074. ELISA plates were blocked, and all sample and secondary antibody dilutions were prepared, in blocking buffer consisting 5% skim milk, 5% bovine serum albumin (BSA) (Fisher Scientific), and 0.1% Tween-20.

Following two washes with PBS-T (PBS with 0.05% Tween-20), the plates were blocked with blocking buffer for 1 h at 37°C. Prior to use, sera were heat-inactivated at 56°C for 30 min and 0.1% Tween-20 was added. Sera were then serially diluted in blocking buffer and added to coated plates in duplicate. Purified recombinant 10-1074 was used to generate standard curves as appropriate. Samples were incubated at 37°C for 1 h, then plates were washed five times with PBS-T. To assess 10-1074 serum concentration, a 1:5000 dilution of a horseradish peroxidase (HRP)-conjugated antibody targeting the IgG Fc (Jackson Immuno Research) was added. For ADA ELISAs, a 1:8000 dilution of an HRP-conjugated anti-human kappa light chain antibody (Millipore Sigma) was used to quantify 10-1074 binding antibodies. Following a 1 h incubation at 37°C, the plates were washed ten times with PBS-T. Plates were developed with 3,3’,5,5’-Tetramethylbenzidine (TMB) Substrate Solution (Thermo Fisher Scientific) for 2-10 min, depending on the assay, and the reaction was stopped with TMB Stop Solution (KPL).

Absorbance at 450 nm was measured using a BioTek Synergy Neo2 plate reader (Agilent). Endpoint titers determined by identifying the serum dilution that produced an optical density (O.D.) of 0.2.

### AAV neutralization assay

AAV neutralization assays were performed as previously described^46,47^. Sera were heat inactivated, diluted 1:5 in DMEM, and then serially diluted four-fold to yield final assay dilutions ranging from 1:10 to 1:5,120. Each serum dilution was tested in duplicate in a 96-well plate and mixed 1:1 (v/v) with 10^10^ vg mL^-1^ of AAV9.CAG.fLuc vector before incubation at 37°C for 30 min. Next, 25,000 HEK293T cells were added to each well. After incubation at 37°C for 24 h, luciferase activity was determined using Britelite Plus (Revvity) and read on a BioTek Synergy Neo2 plate reader (Agilent). The transfer plasmid for pAAV.CAG.fLuc was a gift from Dr. Mark Kay (Addgene # 83281).

### ELISpot assays

MultiScreen-HA Filter plates (MAHA S4519; Milipore) were coated overnight at 4°C with 5 µg/mL mouse anti-human IFN-γ (Pharmingen) in PBS, washed four times with RPMI supplemented with 10% FBS and 1% penicillin/streptomycin (FRPMI), and blocked with FRPMI for 1 h at 37°C. Cryopreserved PBMCs previously isolated by density gradient centrifugation using Ficoll-Pacque PLUS (Cytiva) were thawed, washed in FRPMI with 50 U/mL benzonase, resuspended in FRPMI, and rested for at at 37°C for a minimum of 2 h. Peptide pools were synthesized by GenScript and covered the variable regions of the bNAb heavy (10-1074 VarH) and light (10-1074 VarL) chains; the LS-Furin-P2A region; the IgG1 constant heavy chain region; and the lambda light chain constant regions. Peptide pools, positive-control staphylococcal enterotoxin B (SEB; ToxTech), or DMSO (Sigma) were dispensed into the 96-well plate in duplicate and 200,000–250,000 PBMCs were added per well to yield final concentrations of 10 µg mL⁻¹ for each peptide pool, 4 µg mL⁻¹ for SEB, and 0.1% DMSO for vehicle controls. Following incubation for 20 h at 37°C, plates were washed, and incubated with 1 µg mL^-1^ biotin-conjugated anti-human IFN-γ mAb in PBS-T + 1% FBS (PBS-T-FBS) for 3 h 37°C. Plates were then incubated with a 1:1000 dilution of streptavidin-conjugated alkaline phosphatase (Rockland) in PBS-T-FBS for 1 h at 37 °C, following which they were washed four times with PBS-T. Spots were developed with One-Step NBT/BCIP (Thermo Scientific) and quantified using CTL ImmunoSpot 7.0 software (Cellular Technology Ltd).

### H&E and Immunohistochemistry

Rhesus macaque muscle tissue from the upper left quadericeps was fixed in 10% neutral buffered formalin (NBF), processed, and blocked in paraffin for histological analysis. Samples were sectioned at 5μm and stained with hematoxylin and eosin (H&E) for routine histopathology. Staining for multiple antibodies, including CD3 (1:200, Abcam, ab16669); CD4 (1:200, Abcam, ab13316); CD19 (1:200, Abcam, ab134114); CD68 (1:200, Thermo Fisher, MA5-13324); FoxP3 (1:200, Abcam, ab20034); CD56 (Leica, CD56-504) and PD-1 (1:200, Sino Biologicals, 90305-MM09) was performed on the Bond RX automated system with the Bond Polymer Refine Red Detection (DS9390) (Leica) according to the manufacturer’s instructions. Tissue sections were dewaxed with Bond Dewaxing Solution (Leica) at 72°C for 30 min. Heat-induced epitope retrieval was performed using Epitope Retrieval Solution (Leica), heated to 100°C for 20 min. Tissue sections were visualized by light microscopy on an Olympus BX51 microscope. Photographs were acquired using an Olympus DP73 camera and histopathology was assessed in a blinded fashion by a board-certified veterinary pathologist.

### Luminex assays

Serum cyto/chemokine levels were determined using the NHP XL Cytokine Luminex Performance Premixed 8-plex Kit (R&D Systems, cat# FCSTM21-08) according to the manufacturer’s instructions. Samples were acquired in duplicate on a Bio-Plex 200 System (Bio-Rad Laboratories, Hercules CA) and analyzed using Bio-Plex Manager Software (Bio-Rad).

## Statistical analysis

Data were analyzed using GraphPad Prism v10.4 (GraphPad, La Jolla, CA). Comparisons of groups were performed as indicated in manuscript and/or reported in the figures legends with statistical significance reported as a p value ≤ 0.05.

## Acknowledgements

The authors would like to thank M. Farzan, C.C. Bailey, M.A. Martins, and M.D. Alpert for their discussions and insights regarding this study; the Emory National Primate Research Center (ENPRC) staff for their tremendous help in completing this study; and M.E. Davis-Gardner for her comments and edits to the manuscript. This work was supported in part by National Institutes of Health awards R01AI167724 (M.R.G.) and R01DA056770 (M.R.G.). Additional support was provided from the NIH Office of Research Infrastructure Programs (ORIP) P51OD11132 to ENPRC, U42OD011023 to ENPRC, and P30AI050409 to the Emory University Center for AIDS Research. Next generation sequencing services were provided by the Emory NPRC Genomics Core (RRID:SCR_026418) which is supported in part by NIH P51OD011132. Sequencing data was acquired on an Illumina NovaSeq 6000 funded by NIH S10OD026799.

## Author Contributions

MRG conceived the study and acquired funding. MK, AAK, IL, PD, YB, MY-HL, SL, MC, DM, EDD, and MRG performed experiments. SW, CW, SE, RLS, JSW, and EHC were responsible for performing the required procedures and sample collection of the nonhuman primate study. YN provided critical reagents and input into the study design. JX and GG were responsible for production and quality control of the AAV vectors. MK, MYL, GKT, SL, MC, SV, DAK, INM, SEB, and MRG were responsible for data analysis. MK, SEB, and MRG wrote the initial draft of the manuscript which was reviewed and approved by all coauthors.

## Data, code and material availability

Data supporting the findings appear in the manuscript and are available from the corresponding author.

---

**Figure S1.**
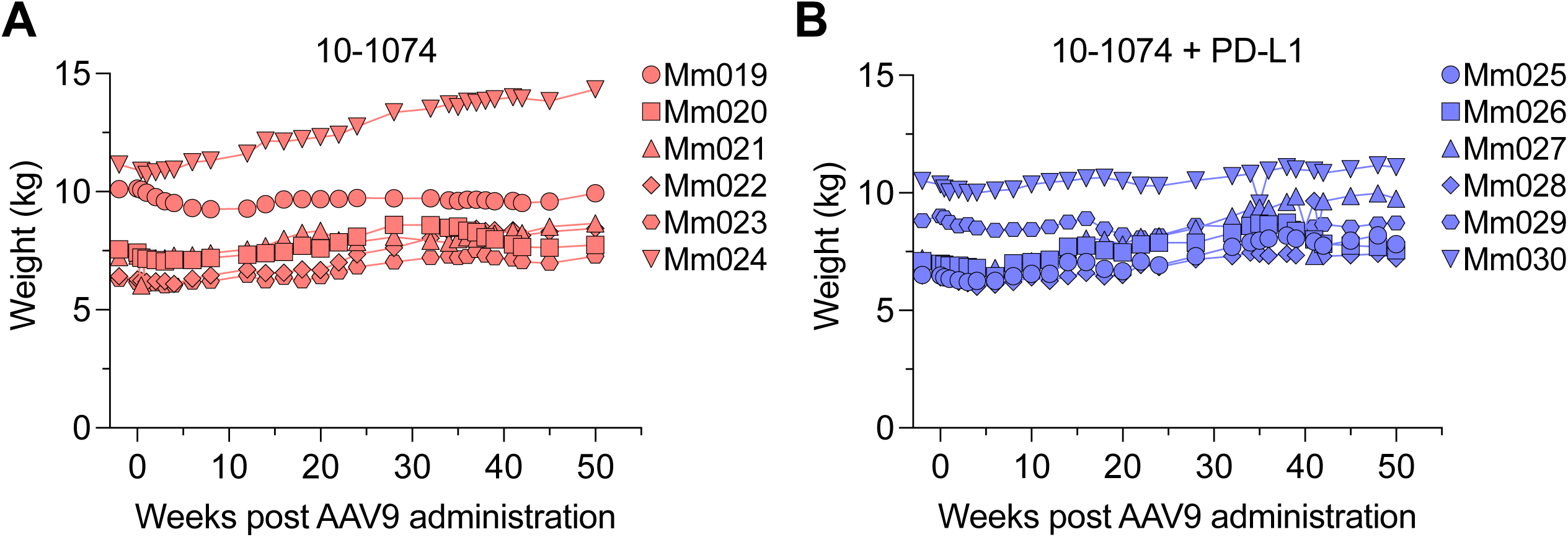
Macaque weight gain over the course of the study. Weight gain over the course of 52 weeks in macaques that received **(A)** AAV9.10-1074 only or **(B)** AAV9.10-1074 plus AAV9.PD-L1.

**Table S1:** Characteristics of the 12 rhesus macaques enrolled in this study.

| **Macaque code** | **Group** | **Age (years)** | **Weight (kg)** | **Sex** | ***Mamu-B\*17*** | ***Mamu-B\*08*** | ***Mamu-A\*01*** | ***Mamu-A\*02*** |
| --- | --- | --- | --- | --- | --- | --- | --- | --- |
| Mm019 | 10-1074-only | 9.7 | 10.11 | F | - | - | + | - |
| Mm020 | 10-1074-only | 3.8 | 7.55 | M | - | - | + | - |
| Mm021 | 10-1074-only | 3.8 | 7.23 | M | - | - | - | - |
| Mm022 | 10-1074-only | 4.0 | 6.42 | M | - | - | - | - |
| Mm023 | 10-1074-only | 3.8 | 6.28 | M | - | - | - | - |
| Mm024 | 10-1074-only | 4.9 | 11.14 | M |  |  |  |  |
| Mm025 | 10-1074 plus PD-L1 | 3.8 | 6.5 | M | - | - | - | - |
| Mm026 | 10-1074 plus PD-L1 | 4.8 | 7.1 | M | - | - | - | - |
| Mm027 | 10-1074 plus PD-L1 | 3.8 | 7.22 | M | - | - | - | - |
| Mm028 | 10-1074 plus PD-L1 | 3.8 | 6.48 | M | - | - | - | + |
| Mm029 | 10-1074 plus PD-L1 | 6.0 | 8.82 | F | - | - | - | - |
| Mm030 | 10-1074 plus PD-L1 | 6.7 | 10.5 | F | - | - | - | - |

## References and Notes

1. Hammer, S.M., Squires, K.E., Hughes, M.D., Grimes, J.M., Demeter, L.M., Currier, J.S., Eron, J.J., Jr., Feinberg, J.E., Balfour, H.H., Jr., Deyton, L.R., Chodakewitz, J.A., et al. (1997). A controlled trial of two nucleoside analogues plus indinavir in persons with human immunodeficiency virus infection and CD4 cell counts of 200 per cubic millimeter or less. AIDS Clinical Trials Group 320 Study Team. N Engl J Med 337, 725–733. 10.1056/nejm199709113371101.

2. Lundgren, J.D., Babiker, A.G., Gordin, F., Emery, S., Grund, B., Sharma, S., Avihingsanon, A., Cooper, D.A., Fätkenheuer, G., Llibre, J.M., Molina, J.M., et al. (2015). Initiation of Antiretroviral Therapy in Early Asymptomatic HIV Infection. N Engl J Med 373, 795–807. 10.1056/NEJMoa1506816.

3. Cohen, M.S., Chen, Y.Q., McCauley, M., Gamble, T., Hosseinipour, M.C., Kumarasamy, N., Hakim, J.G., Kumwenda, J., Grinsztejn, B., Pilotto, J.H., Godbole, S.V., et al. (2016). Antiretroviral Therapy for the Prevention of HIV-1 Transmission. N Engl J Med 375, 830–839. 10.1056/NEJMoa1600693.

4. Swindells, S., Andrade-Villanueva, J.F., Richmond, G.J., Rizzardini, G., Baumgarten, A., Masiá, M., Latiff, G., Pokrovsky, V., Bredeek, F., Smith, G., Cahn, P., et al. (2020). Long-Acting Cabotegravir and Rilpivirine for Maintenance of HIV-1 Suppression. N Engl J Med 382, 1112–1123. 10.1056/NEJMoa1904398.

5. Finzi, D., Blankson, J., Siliciano, J.D., Margolick, J.B., Chadwick, K., Pierson, T., Smith, K., Lisziewicz, J., Lori, F., Flexner, C., Quinn, T.C., et al. (1999). Latent infection of CD4+ T cells provides a mechanism for lifelong persistence of HIV-1, even in patients on effective combination therapy. Nat Med 5, 512–517. 10.1038/8394.

6. Bekker, L.G., Das, M., Abdool Karim, Q., Ahmed, K., Batting, J., Brumskine, W., Gill, K., Harkoo, I., Jaggernath, M., Kigozi, G., Kiwanuka, N., et al. (2024). Twice-Yearly Lenacapavir or Daily F/TAF for HIV Prevention in Cisgender Women. N Engl J Med 391, 1179–1192. 10.1056/NEJMoa2407001.

7. Bar-On, Y., Gruell, H., Schoofs, T., Pai, J.A., Nogueira, L., Butler, A.L., Millard, K., Lehmann, C., Suárez, I., Oliveira, T.Y., Karagounis, T., et al. (2018). Safety and antiviral activity of combination HIV-1 broadly neutralizing antibodies in viremic individuals. Nat Med 24, 1701–1707. 10.1038/s41591-018-0186-4.

8. Mendoza, P., Gruell, H., Nogueira, L., Pai, J.A., Butler, A.L., Millard, K., Lehmann, C., Suárez, I., Oliveira, T.Y., Lorenzi, J.C.C., Cohen, Y.Z., et al. (2018). Combination therapy with anti-HIV-1 antibodies maintains viral suppression. Nature 561, 479–484. 10.1038/s41586-018-0531-2.

9. Corey, L., Gilbert, P.B., Juraska, M., Montefiori, D.C., Morris, L., Karuna, S.T., Edupuganti, S., Mgodi, N.M., deCamp, A.C., Rudnicki, E., Huang, Y., et al. (2021). Two Randomized Trials of Neutralizing Antibodies to Prevent HIV-1 Acquisition. N Engl J Med 384, 1003–1014. 10.1056/NEJMoa2031738.

10. Gaebler, C., Nogueira, L., Stoffel, E., Oliveira, T.Y., Breton, G., Millard, K.G., Turroja, M., Butler, A., Ramos, V., Seaman, M.S., Reeves, J.D., et al. (2022). Prolonged viral suppression with anti-HIV-1 antibody therapy. Nature 606, 368–374. 10.1038/s41586-022-04597-1.

11. Sneller, M.C., Blazkova, J., Justement, J.S., Shi, V., Kennedy, B.D., Gittens, K., Tolstenko, J., McCormack, G., Whitehead, E.J., Schneck, R.F., Proschan, M.A., et al. (2022). Combination anti-HIV antibodies provide sustained virological suppression. Nature 606, 375–381. 10.1038/s41586-022-04797-9.

12. Martinez-Navio, J.M., Fuchs, S.P., Mendes, D.E., Rakasz, E.G., Gao, G., Lifson, J.D., and Desrosiers, R.C. (2020). Long-Term Delivery of an Anti-SIV Monoclonal Antibody With AAV. Front Immunol 11, 449. 10.3389/fimmu.2020.00449.

13. Balazs, A.B., Chen, J., Hong, C.M., Rao, D.S., Yang, L., and Baltimore, D. (2011). Antibody-based protection against HIV infection by vectored immunoprophylaxis. Nature 481, 81–84. 10.1038/nature10660.

14. Balazs, A.B., Ouyang, Y., Hong, C.M., Chen, J., Nguyen, S.M., Rao, D.S., An, D.S., and Baltimore, D. (2014). Vectored immunoprophylaxis protects humanized mice from mucosal HIV transmission. Nat Med 20, 296–300. 10.1038/nm.3471.

15. Gardner, M.R., Kattenhorn, L.M., Kondur, H.R., von Schaewen, M., Dorfman, T., Chiang, J.J., Haworth, K.G., Decker, J.M., Alpert, M.D., Bailey, C.C., Neale, E.S., Jr., et al. (2015). AAV-expressed eCD4-Ig provides durable protection from multiple SHIV challenges. Nature 519, 87–91. 10.1038/nature14264.

16. Welles, H.C., Jennewein, M.F., Mason, R.D., Narpala, S., Wang, L., Cheng, C., Zhang, Y., Todd, J.P., Lifson, J.D., Balazs, A.B., Alter, G., et al. (2018). Vectored delivery of anti-SIV envelope targeting mAb via AAV8 protects rhesus macaques from repeated limiting dose intrarectal swarm SIVsmE660 challenge. PLoS Pathog 14, e1007395. 10.1371/journal.ppat.1007395.

17. Martinez-Navio, J.M., Fuchs, S.P., Pantry, S.N., Lauer, W.A., Duggan, N.N., Keele, B.F., Rakasz, E.G., Gao, G., Lifson, J.D., and Desrosiers, R.C. (2019). Adeno-Associated Virus Delivery of Anti-HIV Monoclonal Antibodies Can Drive Long-Term Virologic Suppression. Immunity 50, 567–575.e565. 10.1016/j.immuni.2019.02.005.

18. Fuchs, S.P., Martinez-Navio, J.M., Piatak, M., Jr., Lifson, J.D., Gao, G., and Desrosiers, R.C. (2015). AAV-Delivered Antibody Mediates Significant Protective Effects against SIVmac239 Challenge in the Absence of Neutralizing Activity. PLoS Pathog 11, e1005090. 10.1371/journal.ppat.1005090.

19. Saunders, K.O., Wang, L., Joyce, M.G., Yang, Z.Y., Balazs, A.B., Cheng, C., Ko, S.Y., Kong, W.P., Rudicell, R.S., Georgiev, I.S., Duan, L., et al. (2015). Broadly Neutralizing Human Immunodeficiency Virus Type 1 Antibody Gene Transfer Protects Nonhuman Primates from Mucosal Simian-Human Immunodeficiency Virus Infection. J Virol 89, 8334–8345. 10.1128/jvi.00908-15.

20. Martinez-Navio, J.M., Fuchs, S.P., Pedreño-López, S., Rakasz, E.G., Gao, G., and Desrosiers, R.C. (2016). Host Anti-antibody Responses Following Adeno-associated Virus-mediated Delivery of Antibodies Against HIV and SIV in Rhesus Monkeys. Mol Ther 24, 76–86. 10.1038/mt.2015.191.

21. Gardner, M.R., Fetzer, I., Kattenhorn, L.M., Davis-Gardner, M.E., Zhou, A.S., Alfant, B., Weber, J.A., Kondur, H.R., Martinez-Navio, J.M., Fuchs, S.P., Desrosiers, R.C., et al. (2019). Anti-drug Antibody Responses Impair Prophylaxis Mediated by AAV-Delivered HIV-1 Broadly Neutralizing Antibodies. Mol Ther 27, 650–660. 10.1016/j.ymthe.2019.01.004.

22. Priddy, F.H., Lewis, D.J.M., Gelderblom, H.C., Hassanin, H., Streatfield, C., LaBranche, C., Hare, J., Cox, J.H., Dally, L., Bendel, D., Montefiori, D., et al. (2019). Adeno-associated virus vectored immunoprophylaxis to prevent HIV in healthy adults: a phase 1 randomised controlled trial. Lancet HIV 6, e230–e239. 10.1016/s2352-3018(19)30003-7.

23. Casazza, J.P., Cale, E.M., Narpala, S., Yamshchikov, G.V., Coates, E.E., Hendel, C.S., Novik, L., Holman, L.A., Widge, A.T., Apte, P., Gordon, I., et al. (2022). Safety and tolerability of AAV8 delivery of a broadly neutralizing antibody in adults living with HIV: a phase 1, dose-escalation trial. Nat Med 28, 1022–1030. 10.1038/s41591-022-01762-x.

24. Burton, D.R., and Hangartner, L. (2016). Broadly Neutralizing Antibodies to HIV and Their Role in Vaccine Design. Annu Rev Immunol 34, 635–659. 10.1146/annurev-immunol-041015-055515.

25. Gernoux, G., Gruntman, A.M., Blackwood, M., Zieger, M., Flotte, T.R., and Mueller, C. (2020). Muscle-Directed Delivery of an AAV1 Vector Leads to Capsid-Specific T Cell Exhaustion in Nonhuman Primates and Humans. Mol Ther 28, 747–757. 10.1016/j.ymthe.2020.01.004.

26. Adriouch, S., Franck, E., Drouot, L., Bonneau, C., Jolinon, N., Salvetti, A., and Boyer, O. (2011). Improved Immunological Tolerance Following Combination Therapy with CTLA-4/Ig and AAV-Mediated PD-L1/2 Muscle Gene Transfer. Front Microbiol 2, 199. 10.3389/fmicb.2011.00199.

27. Xiao, Y., Muhuri, M., Li, S., Qin, W., Xu, G., Luo, L., Li, J., Letizia, A.J., Wang, S.K., Chan, Y.K., Wang, C., et al. (2019). Circumventing cellular immunity by miR142-mediated regulation sufficiently supports rAAV-delivered OVA expression without activating humoral immunity. JCI Insight 5. 10.1172/jci.insight.99052.

28. Ferreira, V., Twisk, J., Kwikkers, K., Aronica, E., Brisson, D., Methot, J., Petry, H., and Gaudet, D. (2014). Immune responses to intramuscular administration of alipogene tiparvovec (AAV1-LPL(S447X)) in a phase II clinical trial of lipoprotein lipase deficiency gene therapy. Hum Gene Ther 25, 180–188. 10.1089/hum.2013.169.

29. Fuchs, S.P., Mondragon, P.G., Zabizhin, R., Tomer, S., Wang, L., Cook, E., Dudley, D.M., Weisgrau, K.L., Furlott, J., Coonen, J., Alexander, E., et al. (2025). Transient rapamycin treatment avoids unwanted host immune responses toward AAV-delivered anti-HIV antibodies. Nat Commun 16, 8906. 10.1038/s41467-025-63970-6.

30. Mendell, J.R., Rodino-Klapac, L.R., Rosales, X.Q., Coley, B.D., Galloway, G., Lewis, S., Malik, V., Shilling, C., Byrne, B.J., Conlon, T., Campbell, K.J., et al. (2010). Sustained alpha-sarcoglycan gene expression after gene transfer in limb-girdle muscular dystrophy, type 2D. Ann Neurol 68, 629–638. 10.1002/ana.22251.

31. Cao, O., Dobrzynski, E., Wang, L., Nayak, S., Mingle, B., Terhorst, C., and Herzog, R.W. (2007). Induction and role of regulatory CD4+CD25+ T cells in tolerance to the transgene product following hepatic in vivo gene transfer. Blood 110, 1132–1140. 10.1182/blood-2007-02-073304.

32. Fuchs, S.P., Martinez-Navio, J.M., Rakasz, E.G., Gao, G., and Desrosiers, R.C. (2020). Liver-Directed but Not Muscle-Directed AAV-Antibody Gene Transfer Limits Humoral Immune Responses in Rhesus Monkeys. Mol Ther Methods Clin Dev 16, 94–102. 10.1016/j.omtm.2019.11.010.

33. Ardeshir, A., O’Hagan, D., Mehta, I., Shandilya, S., Hopkins, L.L.J., Adamson, L., Kuroda, M.J., Hahn, P.A., da Costa, L.A.B., Fuchs, S.P., Martinez-Navio, J.M., et al. (2025). Determinants of successful AAV-vectored delivery of HIV-1 bNAbs in early life. Nature 645, 1020–1028. 10.1038/s41586-025-09330-2.

34. Freeman, G.J., Long, A.J., Iwai, Y., Bourque, K., Chernova, T., Nishimura, H., Fitz, L.J., Malenkovich, N., Okazaki, T., Byrne, M.C., Horton, H.F., et al. (2000). Engagement of the PD-1 immunoinhibitory receptor by a novel B7 family member leads to negative regulation of lymphocyte activation. J Exp Med 192, 1027–1034. 10.1084/jem.192.7.1027.

35. Davis-Gardner, M.E., Weber, J.A., Xie, J., Pekrun, K., Alexander, E.A., Weisgrau, K.L., Furlott, J.R., Rakasz, E.G., Kay, M.A., Gao, G., Farzan, M., et al. (2023). A strategy for high antibody expression with low anti-drug antibodies using AAV9 vectors. Front Immunol 14, 1105617. 10.3389/fimmu.2023.1105617.

36. Gautam, R., Nishimura, Y., Gaughan, N., Gazumyan, A., Schoofs, T., Buckler-White, A., Seaman, M.S., Swihart, B.J., Follmann, D.A., Nussenzweig, M.C., and Martin, M.A. (2018). A single injection of crystallizable fragment domain-modified antibodies elicits durable protection from SHIV infection. Nat Med 24, 610–616. 10.1038/s41591-018-0001-2.

37. McMurphy, T.B., Park, A., Heizer, P.J., Bottenfield, C., Kurasawa, J.H., Ikeda, Y., and Doran, M.R. (2024). AAV-mediated co-expression of an immunogenic transgene plus PD-L1 enables sustained expression through immunological evasion. Sci Rep 14, 28853. 10.1038/s41598-024-75698-2.

38. Kahan, R., Alderete, I.S., Gao, Q., Hughes, B., Zhang, M., Gonzalez, T., Rosales, A., Carney, J., Aykun, N., Abraham, N., Hassan, A., et al. (2025). Adeno-associated virus-mediated transduction of PD-L1 in a rodent lung transplant model. Am J Transplant 25, 2082–2089. 10.1016/j.ajt.2025.05.029.

39. Käyhty, P., Nieminen, T., Eriksson, R.A.E., Tumelius, R., Tawfek, A., Laidinen, S., Ruotsalainen, A.K., Bailey, A., Galibert, L., Lesch, H.P., Ylä-Herttuala, S., et al. (2026). The co-delivery of Programmed Death 1 ligands enhances and prolongs rAAV-mediated gene expression in pre-immunized mice. Gene Ther 33, 127–137. 10.1038/s41434-025-00588-9.

40. Mueller, C., Ratner, D., Zhong, L., Esteves-Sena, M., and Gao, G. (2012). Production and discovery of novel recombinant adeno-associated viral vectors. Curr Protoc Microbiol Chapter 14, Unit14D.11. 10.1002/9780471729259.mc14d01s26.

41. Shingai, M., Donau, O.K., Schmidt, S.D., Gautam, R., Plishka, R.J., Buckler-White, A., Sadjadpour, R., Lee, W.R., LaBranche, C.C., Montefiori, D.C., Mascola, J.R., et al. (2012). Most rhesus macaques infected with the CCR5-tropic SHIV(AD8) generate cross-reactive antibodies that neutralize multiple HIV-1 strains. Proc Natl Acad Sci U S A 109, 19769–19774. 10.1073/pnas.1217443109.

42. Silvestri, G., Sodora, D.L., Koup, R.A., Paiardini, M., O’Neil, S.P., McClure, H.M., Staprans, S.I., and Feinberg, M.B. (2003). Nonpathogenic SIV infection of sooty mangabeys is characterized by limited bystander immunopathology despite chronic high-level viremia. Immunity 18, 441–452. 10.1016/s1074-7613(03)00060-8.

43. Gardner, M.R., Fellinger, C.H., Prasad, N.R., Zhou, A.S., Kondur, H.R., Joshi, V.R., Quinlan, B.D., and Farzan, M. (2016). CD4-Induced Antibodies Promote Association of the HIV-1 Envelope Glycoprotein with CD4-Binding Site Antibodies. J Virol 90, 7822–7832. 10.1128/jvi.00803-16.

44. Fellinger, C.H., Gardner, M.R., Bailey, C.C., and Farzan, M. (2017). Simian Immunodeficiency Virus SIVmac239, but Not SIVmac316, Binds and Utilizes Human CD4 More Efficiently than Rhesus CD4. J Virol 91. 10.1128/jvi.00847-17.

45. Gardner, M.R., Fellinger, C.H., Kattenhorn, L.M., Davis-Gardner, M.E., Weber, J.A., Alfant, B., Zhou, A.S., Prasad, N.R., Kondur, H.R., Newton, W.A., Weisgrau, K.L., et al. (2019). AAV-delivered eCD4-Ig protects rhesus macaques from high-dose SIVmac239 challenges. Sci Transl Med 11. 10.1126/scitranslmed.aau5409.

46. Gardner, M.R., Mendes, D.E., Muniz, C.P., Martinez-Navio, J.M., Fuchs, S.P., Gao, G., and Desrosiers, R.C. (2022). High concordance of ELISA and neutralization assays allows for the detection of antibodies to individual AAV serotypes. Mol Ther Methods Clin Dev 24, 199–206. 10.1016/j.omtm.2022.01.003.

47. Verma, S., Nwosu, S.N., Razdan, R., Upadhyayula, S.R., Phan, H.C., Koroma, A.A., Leguizamo, I., Correa, N.S., Kuipa, M., Lee, D., Vanderford, T.H., et al. (2023). Seroprevalence of Adeno-Associated Virus Neutralizing Antibodies in Males with Duchenne Muscular Dystrophy. Hum Gene Ther 34, 430–438. 10.1089/hum.2022.081.

